# Use of Multiple Pharmacodynamic Measures to Deconstruct the Nix-TB Regimen in a Short-Course Murine Model of Tuberculosis

**DOI:** 10.1101/2023.11.08.566205

**Authors:** M.A. Lyons, A. Obregon-Henao, M.E. Ramey, A.A. Bauman, S. Pauly, K. Rossmassler, J. Reid, B. Karger, N.D. Walter, G.T. Robertson

## Abstract

A major challenge for tuberculosis (TB) drug development is to prioritize promising combination regimens from a large and growing number of possibilities. This includes demonstrating individual drug contributions to the activity of higher-order combinations. A BALB/c mouse TB infection model was used to evaluate the contributions of each drug and pairwise combination in the clinically relevant Nix-TB regimen (bedaquiline-pretomanid-linezolid [BPaL]) during the first three weeks of treatment at human equivalent doses. RS ratio, an exploratory pharmacodynamic (PD) marker of ongoing *Mycobacterium tuberculosis* rRNA synthesis, to-gether with solid culture CFU and liquid culture time to positivity (TTP) were used as PD markers of treatment response in lung tissue; and their time course profiles were mathematically modeled using rate equations with pharmacologically interpretable parameters. Antimicrobial interactions were quantified using Bliss independence and Isserlis formulas. Subadditive (or antagonistic) and additive effects on bacillary load, assessed by CFU and TTP, were found for bedaquiline-pretomanid and linezolid-containing pairs, respectively. In contrast, subadditive and additive effects on rRNA synthesis were found for pretomanid-linezolid and bedaquiline-containing pairs, respectively. Additionally, accurate predictions of the response to BPaL for all three PD markers were made using only the single-drug and pairwise effects together with an assumption of negligible three-way drug interactions. The results represent an experimental and PD modeling approach aimed at reducing combinatorial complexity and improving the cost-effectiveness of *in vivo* systems for preclinical TB regimen development.

---

Pulmonary tuberculosis (TB) is treated using combinations of three or more antimicrobial drugs (higher-order drug combinations) taken daily or intermittently for at least 4-6 months (1, 2). There is a need, however, for shorter and more effective treatment regimens, especially for drug-resistant disease (3, 4). Mouse TB infection models are the primary *in vivo* systems for assessing the efficacy of new TB drugs and regimens (5, 6). They generate experimental data on pathophysiology, pharmacokinetics (PK), pharmacodynamics (PD), and relapse; and can guide allometric scaling and dose selection for early-phase clinical trials (7–11). Bactericidal activity of one or more drugs during early treatment is assessed by equilibrium exposure-response relationships, or by response-time profiles with sampling throughout the study duration (12, 13). The sterilizing or treatment-shortening activity of regimens is typically expressed as time required to prevent relapse in 95% of mice in the relapsing mouse model which treats mice for varying durations and then quantifies microbiologic relapse after a drug holiday (12). These study types provide a basis for prioritizing promising regimens and demonstrating individual drug contributions to the activity of a combination; with the latter being critical to development of regimens that may contain multiple new investigational drugs (14).

While there are several established and novel methods for murine model evaluation of TB drugs and regimens (15, 16), their use for an exhaustive comparison between the hundreds of currently possible (17) higher-order combinations would be prohibitively expensive and time consuming. Additionally, individual drug contributions to a combination are conventionally assessed by a process of single-drug additions and replacement (18, 19), which only partially accounts for antimicrobial interactions. Quantitative methods that assess the effects of higher-order combinations based on their single-drug and pairwise components have been described for several types of antibiotics using *in vitro* systems (20–24). Their extension to *in vivo* systems could reduce combinatorial complexity in regimen-based development, and better account for the combined action of the individual drugs within a combination.

The present study describes an evaluation of a three-drug regimen based on single-drug and pairwise combination effects in a murine TB model. A BALB/c mouse high-dose aerosol subacute TB infection model was used to examine the approach with treatment groups for the possible combinations of bedaquiline (B), pretomanid (Pa), and linezolid (L) administered once daily at human equivalent doses for 21 days. The PD response in lung tissue was assessed by solid culture CFU counts, liquid culture time to positivity (TTP), and the RS ratio^®^, an exploratory marker that indicates the rate of bacterial rRNA synthesis (25). The numerator of the RS ratio is unstable precursor rRNA (pre-rRNA) spacer sequence and the denominator is stable mature rRNA. The time course profiles of each PD marker were mathematically modeled using rate equations with parameters for bactericidal activity, RS ratio turnover time and equilibration, and a conversion factor between CFU and TTP time-kill profiles. The results included a demonstration of individual drug contributions to the activity of the BPaL combination, with antimicrobial interactions that were quantified relative to Bliss independence for each drug pair, and an Isserlis formula for the three-drug combination (20). As the BPaL (Nix-TB) regimen was recently approved for treatment of highly drug-resistant TB (26, 27), the results provide a clinically relevant starting point for an expanding library of single-drug and pairwise measurements from which predictions of novel higher-order combination regimens can be made and prioritized for further nonclinical or early-phase clinical testing.

## RESULTS

### Observed data: PD response-time profiles in BALB/c mice

The time course of PD response for each drug (B, Pa, L), drug pair (BPa, BL, PaL), and the three-drug BPaL combination were assessed by RS ratio, CFU, and TTP, from terminal sampling of lung tissue collected on treatment days 0 (baseline), 2, 4, 7, 11, 14, and 21; and for an untreated (UNTx) control group on days 2, 4, and 7. The observed data counts are summarized in Table S1. Scatter plots of group mean and SD values for each PD marker versus time, are shown in Figure 1. The general pattern of activity for log_10_CFU and TTP showed an inverse relationship, with monophasic decreases in the former matching corresponding increases in the latter; and which varied uniformly across the treatment groups. The RS ratio profiles showed a biphasic response with an initial rapid reduction that slowed toward a plateau by the end of the first week. For PaL, however, there was a slower rate of reduction without a clear inflection point; and for linezolid monotherapy there was a pronounced increase from baseline to day 4 followed by a variable but downward trend.

**Figure 1:**
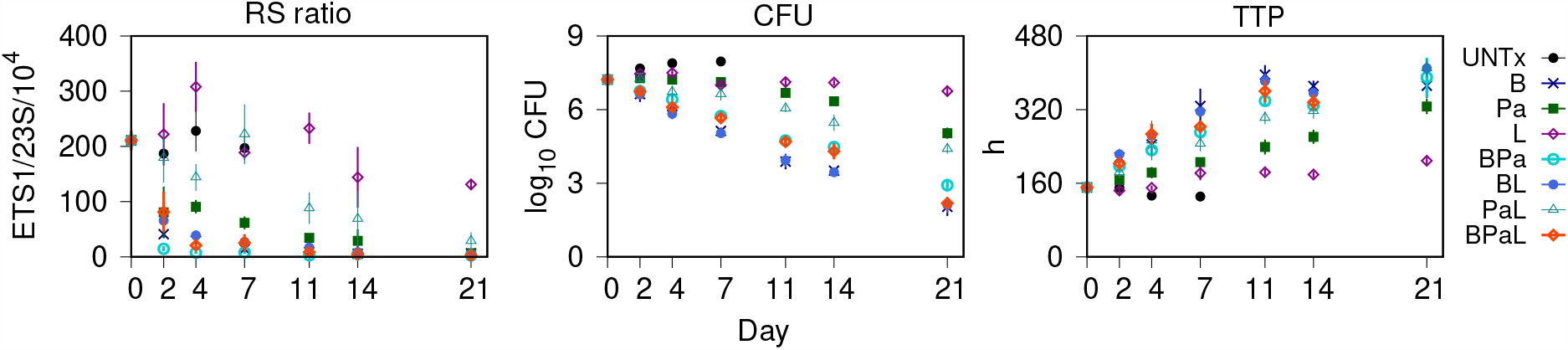
Combined treatment group response profiles of *M. tuberculosis* RS ratio, solid culture CFU, and liquid culture time to positivity (TTP) from BALB/c mouse lungs. Data are the group mean (points) and SD (error bars) of the observed PD response versus treatment day. UNTx, untreated; B, bedaquiline; Pa, pretomanid; L, linezolid; BPa, bedaquiline-pretomanid; BL, bedaquiline-linezolid; PaL, pretomanid-linezolid; and BPaL, bedaquiline-pretomanid-linezolid.

### PD modeling: characterization of drug activity

The PD response profiles were parameterized using rate equations based on synthesis and degradation of pre-rRNA spacer sequences assessed by RS ratio, and bacterial population growth and drug killing effect assessed by CFU and TTP. The model parameters were estimated as probability distributions conditioned on the baseline-normalized group mean values of each PD marker, and are summarized by their distribution mean and SD in Table 1. The rate and extent of drug effect on RS ratio were characterized by a degradation half-life, 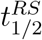, and an equilibration value, *RS*_*ss*_, respectively. The half-life characterizes the initial treatment effect and depends inversely on the degradation rate, while the equilibration value represents the long term equilibrium state. Bactericidal activity was characterized by the drug kill rate constant, *k*_*drug*_, which represents rate of log_10_CFU/lung decrease or TTP increase; together with a solid-liquid culture conversion factor, *τ*, where the liquid culture was a mycobacteria growth indicator tube (MGIT) system. For a particular liquid culture inoculum with matching solid culture CFU, smaller and larger values of *τ* represent, respectively, larger and smaller rates of oxygen consumption yielding corresponding smaller and larger TTP values. The mean (SD) values for the bacterial population growth rate constant, *µ* = 0.72 (0.043)/d, and carrying capacity, *K* = 9.34 × 10^7^ (5.83 × 10^6^) CFU/lung in the absence of drug were determined from the untreated group data. Model simulations of the baseline-normalized PD profiles as fractional effects, *E/E*_0_, are shown together with the observed data in Figure 2; where *E* = *E*(*t*) is the PD response at elapsed time, *t*, from the first dose, and *E*_0_ = *E*(*t* = 0) is the baseline value. The fractional effect for TTP was defined using the reciprocal values, TTP^−1^.

**Table 1:** Pharmacodynamic (PD) model parameter values for bactericidal activity and RS ratio turnover for each treatment group.

| Group | Parameter (units) |  |  |  |
| --- | --- | --- | --- | --- |
| | $k_{drug}$ (1/d) | $\tau$ (h) | $t_{1/2}^{RS}$ (d) | $RS_{ss}$ (ETS1/23S/10<sup>4</sup>) |
| B | 1.29 (0.02) | 28.7 (1.2) | 1.4 (0.1) | 6.1 (1.0) |
| Pa | 0.90 (0.02) | 37.5 (3.5) | 3.5 (0.3) | 7.4 (1.7) |
| L | 0.71 (0.03) | 38.0 (7.7) | 13 (2.4) | 65 (23) |
| BPa | 1.18 (0.01) | 29.6 (1.3) | 0.69 (0.06) | 3.7 (0.6) |
| BL | 1.28 (0.01) | 29.8 (1.3) | 2.1 (0.2) | 6.0 (1.2) |
| PaL | 0.99 (0.02) | 44.0 (2.9) | 6.5 (0.58) | 18 (5) |
| BPaL | 1.25 (0.01) | 30.2 (1.3) | 1.8 (0.1) | 3.7 (0.7) |

Data are posterior distribution mean (SD), sample size equal to 10,000.
$k_{drug}$ , drug kill rate constant; $\tau$ , solid-liquid culture time constant;
$t_{1/2}^{RS}$ , RS ratio half-life; $RS_{ss}$ , RS ratio equilibration constant;
PaL, pretomanid-linezolid; BPaL, bedaquiline-pretomanid-linezolid.

**Figure 2:**
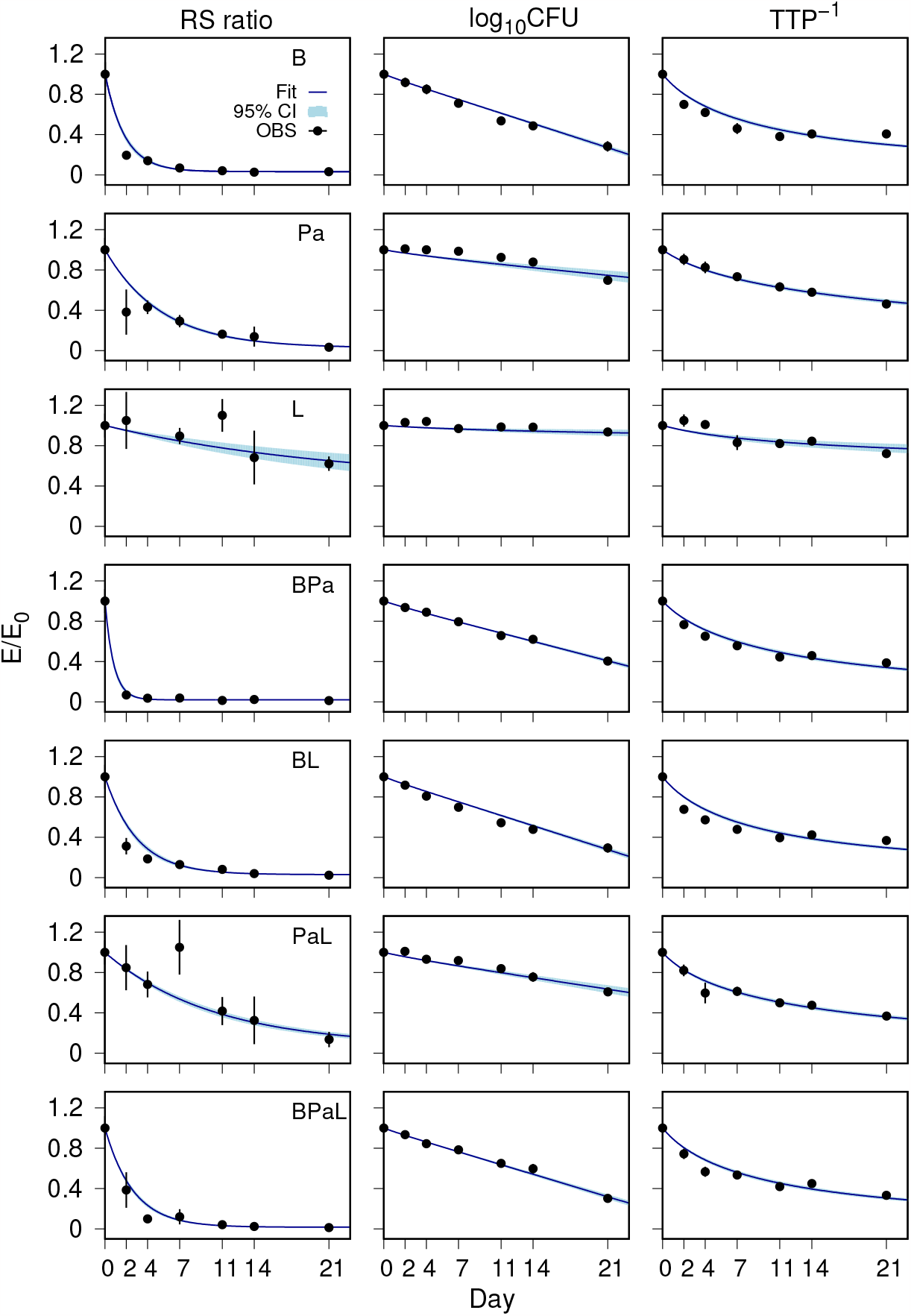
Fractional effect (*E/E*_0_) profiles of *M. tuberculosis* RS ratio, solid culture CFU (log_10_CFU), and reciprocal of the liquid culture time to positivity (TTP^−1^) from BALB/c mouse lungs. Data are model simulations of the predicted mean (solid line with shaded 95% confidence interval) together with observed group mean (points) and SD (error bars) for each treatment group versus time. The observed day 4 RS ratio mean (SD) value not shown in the plot was equal to 1.46 (0.25). B, bedaquiline; Pa, pretomanid; L, linezolid; BPa, bedaquiline-pretomanid; BL, bedaquiline-linezolid; PaL, pretomanid-linezolid; and BPaL, bedaquiline-pretomanid-linezolid.

As single drugs, the effects of B, Pa, and L were distinguished by differences in their PD parameter values and by their profile shapes, with bedaquiline monotherapy showing the largest kill rate, and the most rapid and extensive reduction in RS ratio. While the kill rate constants and overall time course for decline in bacillary burden in the lungs were similar for B, BPa, BL, and BPaL, they show pronounced differences in their effect on RS ratio degradation rate and equilibration values, with BPaL showing the smallest equilibration value. The most rapid effect on RS ratio was observed for mice treated with BPa, which was antagonistic by CFU and TTP measures relative to bedaquiline given as monotherapy. Overall, the RS ratio parameter values and time course profiles exhibited more clearly defined differences between the various tested regimens that were less apparent based on CFU and TTP kill curves alone.

While the PD model assumptions provided an explanatory representation of basic processes and features of the observed profiles for each PD marker, the relatively small but visually apparent errors, or differences between model simulation and observed data, in the first week of treatment indicate transitory drug effect processes that were not accounted for in the model equations. While not especially large, the deviations between model output and observation indicate delayed onset in bactericidal activity in the first week CFU data for Pa, L, and PaL. Similarly, differences between modeled and observed RS ratios for B, BL, and BPaL on day 2 and day 4 indicates a possible additional early phase synthesis-degradation process for bedaquiline exposure. The largest deviation from the modeling assumptions is seen for the RS ratio response to linezolid monotherpy, with the first week increase from baseline indicating a delayed onset of degradation. The errors in the first week of treatment for TTP indicate possible limits on the assumption of similarity between the viable bacilli in the two culture media. After the first week of treatment, however, the modeling errors are smaller and less informative for drug effect processes that deviate from the modeling assumptions. This suggests attainment of equilibrium in drug effects after the first week with response profiles accounted for by simple assumptions of first-order drug kill rates, equivalence between bactericidal activity assessed by log_10_CFU and TTP, and a balance between synthesis and degradation of precursor rRNA.

### PD interactions

Time course profiles of the PD interactions were determined using the fractional effect profiles for RS ratio, log_10_CFU, and TTP^−1^. Figure 3 shows the observed and modeled PD marker data for each drug pair and the pointwise product of the modeled single-drug profiles, where the latter corresponds to an additive effect relative to Bliss independence (statistical independence). Also shown are the interactions calculated as the difference between the two PD response curves, or deviation from additivity, together with the corresponding observed data. Positive deviations represent subadditive (or antagonistic) interactions, resulting from a smaller effect observed for the drug pair than expected by the product of their single drug profiles. Subadditive interactions were found for the BPa CFU and TTP profiles, and for PaL in the RS ratio. All other pairwise effects were found to be approximately additive, without substantive differences between the observed values and those predicted from an independent product assumption. For BPa, the interactions exhibited an initial increasing and variable response that became stable by day 11. Figure 4 shows the observed BPaL data and PD model fits together with the model predicted Isserlis and Bliss formulas. Relative to Bliss independence the combination was found to be subadditive with respect to CFU and TTP, and except for the day 2 value, additive with respect to RS ratio. In contrast, and again with the exception of the day 2 RS ratio value, there were no significant three-way interactions for BPaL when the mutual interactions were accounted for by the Isserlis formula. The absence of three-way interactions used as an assumption (20) provided accurate predictions of BPaL response for all three PD markers based on the responses of the single-drug and pairwise components.

**Figure 3:**
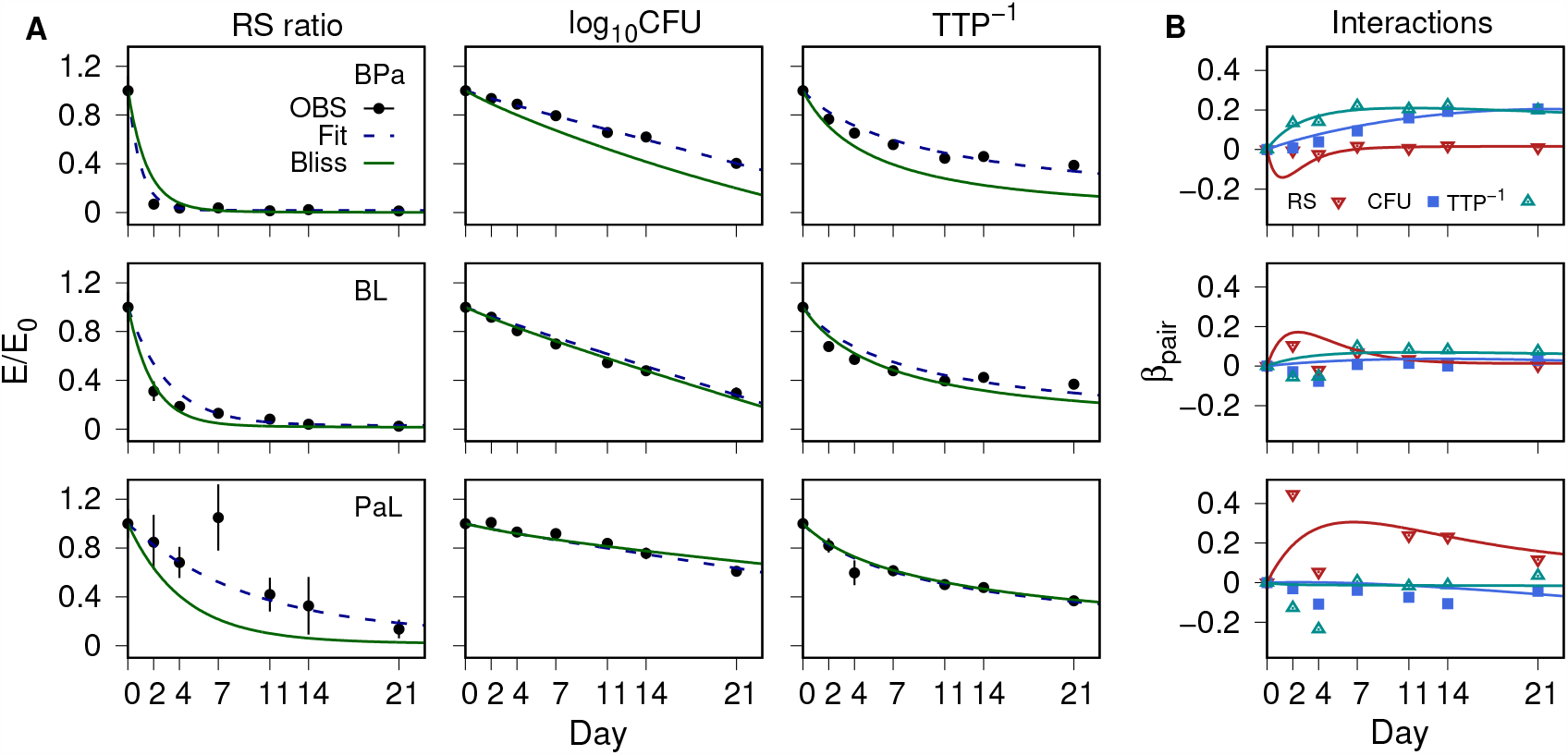
Pairwise PD response profiles and interactions as a deviation from Bliss independence. A PD marker fractional effect (*E/E*_0_), for bedaquiline-pretomanid (BPa), bedaquiline-linezolid (BL) and pretomanid-linezolid (PaL) combinations. Observed data (OBS) group mean (points) and SD (error bars), together with model simulations for the pair (dashed lines) and Bliss independence as pointwise products of single-drug model fits. B Interactions as deviations from Bliss independence. Values are shown for each PD marker for the observed mean values (point symbols) and model predictions (lines).

**Figure 4:**
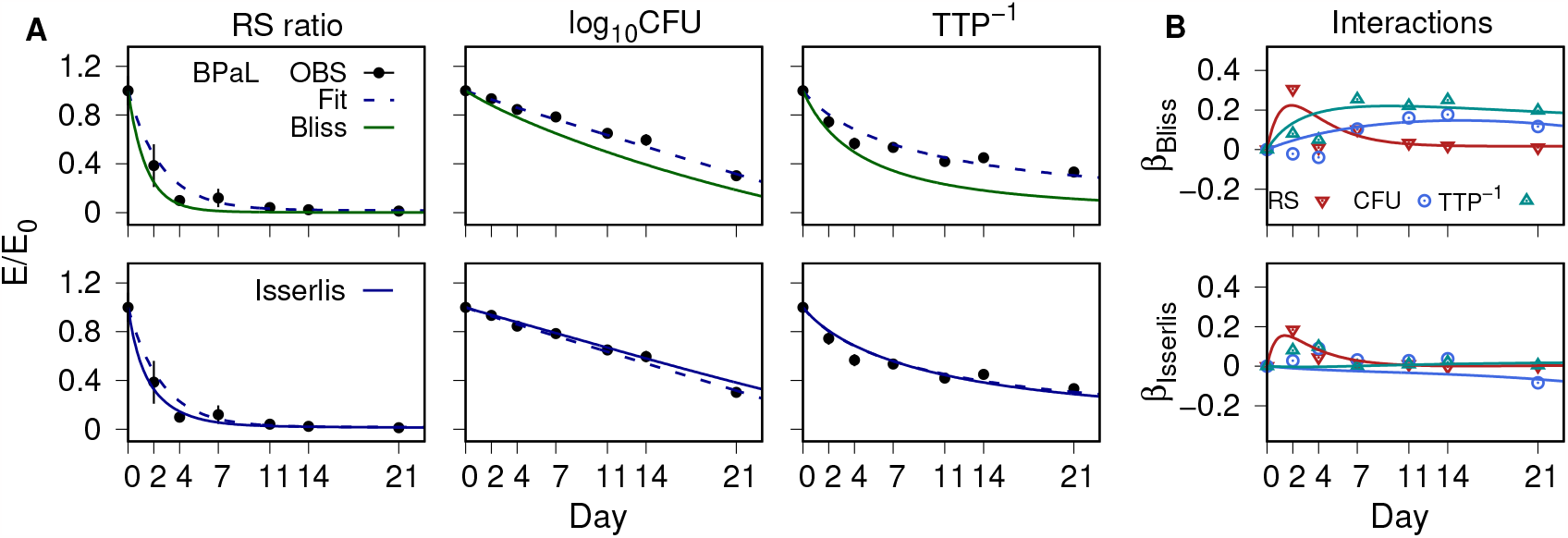
Bedaquiline-pretomanid-linezolid (BPaL) PD response profiles and interactions as deviations from Bliss and Isserlis formulas. A PD marker fractional effect (*E/E*_0_), for bedaquiline-pretomanid-linezolid (BPaL). Observed data (OBS) group mean (points) and SD (error bars), together with model simulations for the combination (dashed lines) and Bliss independence (top row) and Isserlis formula predictions (bottom row). B Interactions as deviations from Bliss independence (top row) and Isserlis formula (bottom row). Values are shown for each PD marker for the observed mean values (point symbols) and model predictions (lines).

## DISCUSSION

While murine models provide critical assessment of new TB drugs and treatment regimens leading up to clinical testing, they can be costly and labor-intensive, which limits their use as high-throughput screens for the large numbers of possible higher-order combination regimens generated from the current drug pipeline (28, 29). The purpose of this study was to examine the use of single-drug and pairwise combinations as a reduced set of targeted drug effect measurements to predict the outcomes of higher-order drug combinations and to demonstrate the component drug contributions to the PD responses of the regimen as a whole. A standard mouse TB infection model with three different PD markers, and the BPaL regimen, were used as a test case. The most common methods for predicting higher-order antibiotic combination effects are based on *in vitro* equilibrium studies measuring bacterial population growth under subinhibitory drug concentrations (20, 21, 23, 30). Here, these methods were adapted to an *in vivo* system with PD response-time profiles and bactericidal concentrations. The results included accurate predictions of BPaL effects on the kinetic profiles of the three PD markers, the characterization of BPaL component drug contributions to the combination based on a minimal set of pharmacologically interpretable parameters, and quantitative measures of drug interactions using two different models for additivity. These results show a potential for single-drug and pairwise measurements to predict higher-order combination effects as an experimentally feasible approach to prioritizing large numbers of novel TB regimens for further nonclinical and early-phase clinical testing; including combinations that would otherwise remain untested.

Demonstration of the individual drug contributions to the activity of a combination is a basic task for development of novel TB regimens (14). For mouse TB models, such demonstration is typically conducted one drug at a time, where efficacy of the combination is assessed with and without each tested drug (18, 19, 31). Separate from investigation of it’s predictive capacity, the Isserlis formula provided a deconstruction of the PD effects of BPaL as a sum of effects arising from single-drug and pairwise products. The PD effects were also characterized for each drug component by a kill rate constant for bactericidal activity, and RS ratio half-life and equilibration as a measure of rRNA synthesis (16, 25). The interactions were quantified over time as deviations from Bliss and Isserlis formulas, with the latter including mutual interactions in contrast to the former which did not. Pretomanid was the only substantively interacting drug, with subadditive effects on bacillary load in combination with bedaquiline, and on rRNA synthesis in combination with linezolid. There were no substantive interactions among the three PD markers for the BL pair, nor any that emerged from all three drugs acting together when mutual interactions were accounted for. The different characterizations of BPaL as subadditive by Bliss independence and additive by the Isserlis formula illustrates the dependence of such classification on how combined action without interaction is defined. The difference in this case is illustrated by the predicitive accuracy of the Isserlis formula compared to Bliss independence. Additionally, such interactions were found to differ depending on the type of PD response, which shows the utility of including multiple PD markers, in this case for rRNA synthesis using the RS ratio and burden using CFU and TTP.

Beyond the methodological limitations in data handling, modeling assumptions, and the sensitivity to variability and error, the main limitation to generalizing the results of this study is the narrow scope, with only one drug combination, one dosage regimen, and one *M. tuberculosis* strain. Evaluation of a wider range of drug classes, including dose ranging for at least the single drug measurements, would better establish a basic pattern of outcomes from this experimental system. Application of the Isserlis formula for prediction required an assumption of negligible higher-order interactions, and was based on previous observation of such effects from *in vitro* antibiotic combination studies including antimycobacterial drugs (20, 21, 23), and from theoretical considerations of locality (32) that might apply to drug-mediated interactions in biochemical processes. While the accuracy of the BPaL results across disparate PD markers may indicate a more general application, there are examples of higher-order or emergent effects that challenge this assumption (22, 30). Another limitation is the absence of selection criteria beyond that of maximum observed effect, where the meaning of interactions with respect to drug resistance and sterilizing activity (18) may require mechanistic understanding and longer treatment duration to assess. Application of this analysis to other PD markers such as lipoarabinomannan (LAM) (33) and 16S rRNA (34) remains untested.

This study combined *in vivo* measurements of drug effect with mathematical modeling as a potential method of prioritizing novel TB regimens for further nonclinical and clinical evaluation. The experimental design was similar to phase 2a early bactericidal activity (EBA) studies (35), and could be further combined with similarly designed *in vitro* time-kill studies to identify translatable elements across the preclinical and early-phase clinical development stages. Consideration of the planned evaluation of novel TB regimens by the

UNITE4TB consortium (36) provides an example of how the approach developed here could be applied. There are eleven planned phase 2b/c studies of mostly 4-drug combinations assembled from eight individual drugs assessed in phase 2a monotherapy testing. In total, 19 phase 2 studies studies will yield an evaluation of 11 combination regimens. Using Isserlis formulas based on single-drug and pairwise outcomes (20), an alternative approach could first measure the 8 single-drug and 28 pairwise combinations of the 8 drugs, which would then yield predictions for 56 three-drug, 70 four-drug, and 56 five-drug combinations. A ranking of promising regimens could be made from comparisons among these predicted outcomes and provide a basis for additional nonclinical testing. Such an approach could provide a more thorough and systematic prioritization of combination regimens for early-phase clinical trials, and the flexibility to incorporate new drugs with minimal experimental testing.

## MATERIALS AND METHODS

### BALB/c mouse high-dose aerosol infection model

All animal studies were performed at Colorado State University in a certified animal biosafety level III facility, and conducted in accordance with guidelines of the Colorado State University Institutional Animal Care and Use Committee (Reference number: 1212). Six- to eight-week-old female pathogen-free BALB/c mice (Jackson Laboratories) were exposed to high-dose aerosol (Glas-Col) of *M. tuberculosis* Erdman from broth culture (optical density at 600nm of ∼0.8) to achieve deposition of ∼4.0 log_10_CFU in the lungs of each mouse (37). Mice were sacrificed for lung

CFU counts one day after infection (day -10) and at day 0 to determine the number of CFU implanted in lungs and the number of bacilli present at the start of treatment, respectively. Antibiotic treatment. Mice were randomized to the different treatment groups. Treatments were initiated 11 days post aerosol (Day 0) and were given once daily by oral gavage, 7 days per week, for the duration of the study. Bedaquiline fumarate was administered at 25 mg/kg. Linezolid was administered at 100 mg/kg. Pretomanid was administered at 50 mg/kg. Bedaquiline fumarate was formulated in 20% hydroxypropyl-b-cyclodextrin solution acidified with 1.5% 1N HCl. Pretomanid was prepared in the CM-2 formulation as previously described (37). Linezolid was prepared in 5% PEG-200/95% [0.5% w:v] methyl-cellulose. In cases of drug combinations, bedaquiline was administered first, followed 1 hr later by pretomanid and/or 4 hrs later by linezolid. Each drug was administered in 0.2 mL volume.

Evaluation of drug activity. Drug bactericidal activity was assessed by lung CFU reductions and time to positivity (TTP) increases, and by RS ratio turnover. Groups of 5 mice each were humanely euthanized 1-day following 2, 4, 7, 11, 14, or 21 consecutive days of drug treatment. At each time point, lung lobes were removed aseptically. The superior and middle lobes (roughly one-third of the lung by mass) were immediately flash frozen under liquid nitrogen for RNA preservation. The remaining two-thirds of the lung was used for CFU and TTP assessments. Details on RS ratio assessement were described previously (25). For CFU plating and TTP assessments, previously frozen lungs were homogenized using the Bertin Precellys CKMix50-7 mL lysis kit (in 4.5 mL PBS with 10% bovine serum albumin (BSA). Lung homogenates were plated in serial dilutions on 0.4% charcoal-supplemented 7H11 agar supplemented with 10% oleic acid, BSA, dextrose, and catalase (OADC) and with selective antibiotics, including cycloheximide (10 mg/L), carbenicillin (50 mg/L).

For TTP assessments, lung homogenates were diluted five-fold in 7H9 media supplemented with 10% OADC and 0.05% Tween80, placed in a 2ml screw-cap tube containing sterile 2mm beads and bead-beat at 2,800 rpm for 10 sec. Homogenates were then inoculated into 7ml mycobacterial growth indicator tubes (MGIT) containing 7H9 media supplemented with 10% OADC, 0.05% Tween80, 0.4% charcoal and BD MGIT PANTA™ Antibiotic Mixture (polymyxin B, amphotericin B, nalidixic acid, trimethoprim and azlocillin), per the manufacturers instructions. Finally, MGIT tubes were incubated in the BD BACTEC MGIT 320 system for at least 42 days at 37°C. TTP was recorded as the time (in hr) for cultures to turn positive. Cultures that were assayed in the MGIT system but remained negative up to day 42 (the accepted cutoff) were recorded as negative.

Pharmacodynamic (PD) modeling and antimicrobial interactions. The PD response profiles for each treatment group were modeled as functions of elapsed time, *t*, from the start of treatment using PD rate equations (38, 39). The PD response for RS ratio was represented by a turnover equation;

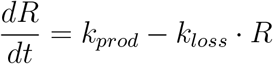

where *R* = *R*(*t*) is the RS ratio with initial condition, *R*(0) = *R*_0_. The rate constants forapparent production, *k*_*prod*_, and fractional loss, *k*_*loss*_, were expressed as a half-life, 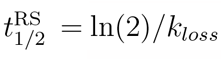, and an equilibration (or steady state) value, *RS*_*ss*_ = *k*_*prod*_*/k*_*loss*_. The CFU and TTP profiles were modeled using logistic growth with constant drug killing effect as,

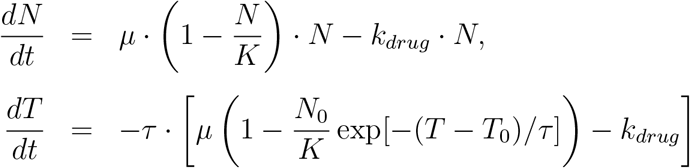

where *N* = *N* (*t*) represents the CFU/lung and *T* = *T* (*t*) is the TTP. The initial conditions were specified as *N* (0) = *N*_0_ and *T* (0) = *T*_0_. The parameter, *µ*, is the bacterial growth rate constant; *K*, the carrying capacity; *k*_*drug*_, the drug kill rate constant; and *τ*, a solid-liquid culture conversion factor between CFU and TTP, with a detailed derivation previously described (40). The model parameters were estimated from the observed data using Bayes’ theorem and Markov chain Monte Carlo (MCMC) simulation (41). The MCMC simulations consisted of ten independent sampling chains of 100,000 iterations each; with every tenth iteration of the last 10,000 retained and aggregated into a final 10,000 iteration sample.

The PD interactions were characterized as functions of time using the fractional response (or survival fraction), *u* = *u*(*t*), for each drug alone, *u*_*i*_, the pairs, *u*_*ij*_, and the three-drug combination, *u*_*ijk*_, where *i, j, k* are drug labels. Pairwise interactions were quantified as the deviation from Bliss independence, *β*_*ij*_ = *u*_*ij*_ − *u*_*i*_*u*_*j*_, and three-way interactions as deviations from an Isserlis formula (20), *β*_*ijk*_ = *u*_*ijk*_ − (*u*_*i*_*u*_*jk*_ + *u*_*j*_*u*_*ki*_ + *u*_*k*_*u*_*ij*_ − 2*u*_*i*_*u*_*j*_*u*_*k*_).

In the absence of pairwise interactions the Isserlis formula reduces to a Bliss independence formula with deviations *β*_*ijk*_ = *u*_*ijk*_ − *u*_*i*_*u*_*j*_*u*_*k*_. Interactions were specified relative to zero deviation (additive) as subadditive (or antagonistic) for positive deviation, or superadditive (or synergistic) for negative deviation.

### Data Handling and Limitations

The observed mouse efficacy data were obtained as terminal samples, which precluded individual PD response-time profiles. The PD profiles used for model parameter estimates were constructed as population summaries from the group mean values calculated with missing data ignored. Negative cultures were treated as missing. The mathematical models were constructed as simple representations of the basic biological processes underlying the observed PD responses during early treatment. Additional features or distinguishing factors related to drug mechanisms of action, such as delays in onset of activity, multiple subpopulations of bacilli, or features that may become evident with longer-duration treatments such as a possibly vanishing RS ratio, were not included in the PD modeling assumptions. The relationship between CFU and TTP included an assumption of similarity between the viable bacillary populations that are detected in the two different culture media, and that this relationship did not change over time. The Isserlis and Bliss formulas are sensitive to measurement uncertainty and variability due to multiplication of error (42).

Software. Model simulations and data analysis were implemented using MCSim Modeling and Simulation Suite (41) (version 6.2.0; [http://www.gnu.org/software/mcsim]), the R statistical software (version 3.3.3; R Development Team, [https://www.R-project.org]) with the CODA package (version 0.18; [https://cran.r-project.org/web/packages/coda]), and gnuplot (Version 5.0. [http://www.gnuplot.info/]). The operating system was Linux (version 3.16.0-4-amd64; Debian distribution [https://www.debian.org]).

## Data Availability

The individual mouse CFU, TTP, and RS ratio data, and computer model simulation files are provided in the Supplementary Material.

## ACKNOWLEDGMENTS

We thank Anne Lenaerts (formerly Colorado State University) for discussions on the study design; Martin Voskuil (University of Colorado Anschutz Medical Campus, CU), Matthew Reichlen (CU), David Hermann (Bill and Melinda Gates Foundation, BMGF), Debra Hanna (BMGF), and Paolo Denti (University of Cape Town, South Africa) for discussions of the results; and Thabo Mabuka (TASK Applied Science, South Africa) for comments and suggestions on the manuscript. We acknowledge the staff of the Laboratory Animal Resources at Colorado State University for their animal care.

HHS | National Institutes of Health (NIH): National Institute of Allergy and Infectious Diseases (NIAID) provided funding to M.A. Lyons under grant number R01AI125454. G.T. Robertson acknowledges support by the Bill and Melinda Gates Foundation under grant number INV-009105. N. D. Walter acknowledges support from Veterans Affairs Merit Award 1I01BX004527-01A1.

NWD and GTR are listed as co-inventors on US patent No. 16/632,310 that pertains to the RS ratio. The remaining authors have no conflicts of interest to declare.

