## Supplemental File for "Use of Multiple Pharmacodynamic Measures to Deconstruct the Nix-TB Regimen in a Short-Course Murine Model of Tuberculosis"

Table S1: Observed data summary and counts.

Figure S1: Untreated group data and model simulation.

Observed data and MCSim files: `Supplementary_Files_Lyons_etal_BPaL_BALBc.zip`

Table S1: Observed data summary for RS ratio, CFU, and time to positivity (TTP).

| Treatment | n | RS ratio | CFU | TTP |
|---|---|---|---|---|
| Pre-treatment | 7 | 7 (0) | 7 (0) | 7 (0) |
| UNTx | 15 | 15 (0) | 15 (0) | 15 (0) |
| B | 30 | 30 (0) | 30 (1) | 30 (3) |
| Pa | 30 | 30 (0) | 30 (0) | 30 (0) |
| L | 30 | 30 (0) | 30 (0) | 30 (0) |
| BPa | 30 | 30 (0) | 30 (0) | 30 (0) |
| BL | 30 | 30 (0) | 30 (0) | 30 (2) |
| PaL | 30 | 30 (0) | 30 (0) | 30 (0) |
| BPaL | 30 | 30 (0) | 30 (0) | 30 (4) |

Data counts, total (culture negative)
n, no. of mice.
UNTx, Untreated; B, bedaquiline; Pa, pretomanid;
L linezolid; BPa, bedaquiline-pretomanid;
BL bedaquiline-linezolid; PaL, pretomanid-linezolid;
BPaL, bedaquiline-pretomanid-linezolid

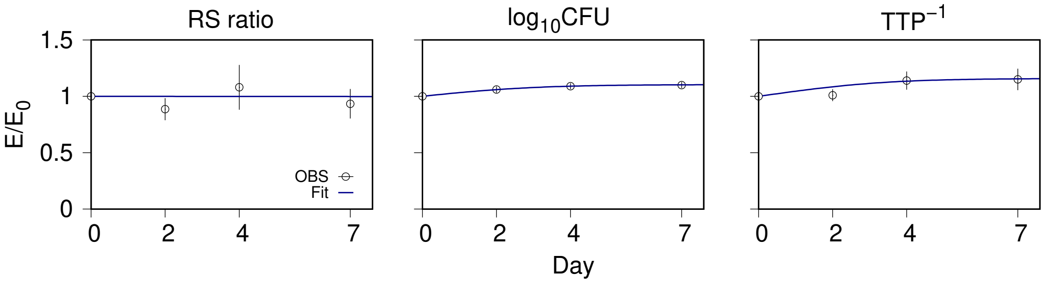


Figure S1: Untreated group data. Fractional effect ($E/E_0$) for the RS ratio, $\log_{10}$CFU, and reciprocal of the time to positivity ($TTP^{-1}$) versus treatment day. Model simulations of logistic growth (solid lines) with growth rate constant $\mu = 0.72$/d and carrying capacity $K = 9.34 \times 10^7$ CFU/lung, together with the observed (OBS) group mean (points) and SD (error bars) The baseline ($E_0$) values (mean [SD], n=7 mice) were 211 (17.8) (ETS1/23S/$10^4$) for RS ratio, 7.22 (0.136) $\log_{10}$ CFU/lung, and 151 (3.26) h for TTP. .

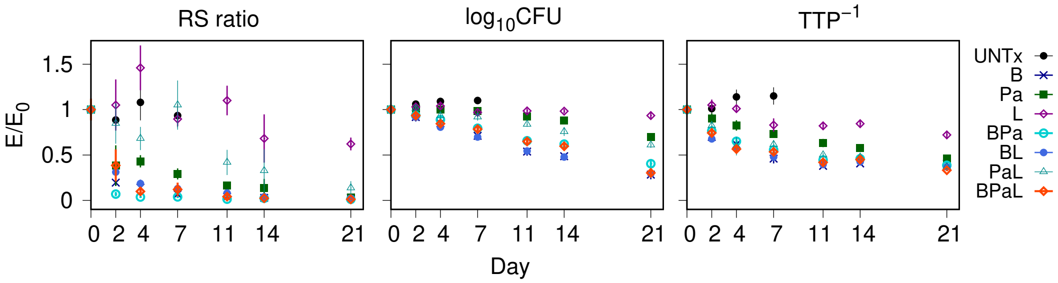

RS ratio
log10CFU
TTP−1
E/E0
1.5
1
0.5
0
0 2 4 7 11 14 21
Day
UNTx
B
Pa
L
BPa
BL
PaL
BPaL


Figure S2:
